# Quantitative live-imaging of *Aquilegia* floral meristems reveals distinct patterns of floral organ initiation and cell-level dynamics of floral meristem termination

**DOI:** 10.1101/2021.10.06.463406

**Authors:** Ya Min, Stephanie J. Conway, Elena M. Kramer

## Abstract

In-depth investigation of any developmental process in plants requires knowledge of both the underpinning molecular networks and how they directly determine patterns of cell division and expansion over time. Floral meristems (FM) produce floral organs, after which they undergo floral meristem termination (FMT), and precise control of organ initiation and FMT is crucial to reproductive success of any flowering plant. Using a live confocal imaging, we characterized developmental dynamics during floral organ primordia initiation and FMT in *Aquilegia coerulea* (Ranunculaceae). Our results have uncovered distinct patterns of primordium initiation between stamens and staminodes compared to carpels, and provided insight into the process of FMT, which is discernable based on cell division dynamics preceding carpel initiation. To our knowledge, this is the first quantitative live imaging of meristem development in a system with numerous whorls of floral organs as well as an apocarpous gynoecium. This study provides crucial information for our understanding of how the spatial-temporal regulation of floral meristem behavior is achieved in both an evolutionary and developmental context.

## INTRODUCTION

The spatial-temporal regulation of cell proliferation and expansion is a fundamental aspect of development in all multicellular organisms. It is particularly crucial in plants because programmed cell death and cell migration play no role in most developmental processes, while morphogenesis and organogenesis occur throughout the entire lifespan of a plant (Steeves & Sussex, 1989). The foundation of continuous growth in a plant is the presence of meristems, which are groups of cells that possess stem cell properties and are typically located at the tips of all growing axes. Meristem activities must be well regulated to maintain a pool of pluripotent cells while also giving rise to differentiating cells. Based on the types of tissues and organs produced, meristems can be categorized into several types. For instance, vegetative meristems produce vegetative organs such as leaves; root apical meristems are responsible for the growth of roots; and in flowering plants, there are inflorescence meristems, which give rise to reproductive branches, and floral meristems (FMs), which produce floral organs. Meristems that produce leaves and roots are indeterminate by nature and can make new organs and tissues continuously. The properties of the inflorescence meristems are more variable, and whether they grow indeterminately or determinately differs across plant lineages (Kirchoff & Claßen-Bockhoff, 2013; Bartlett & Thompson, 2014). FMs, however, are always determinant because every FM is responsible for the production of one flower, and every flower terminates after the production of a relatively finite number of organs, typically culminating in carpels. These consistently determinate meristems are particularly interesting to study in terms of the spatial-temporal regulation of cell behaviors. On the one hand, just like other meristems, FMs need to maintain homeostasis of their stem cell pools to ensure the successive production of organs. On the other hand, this proliferation must be terminated at a specific time point during floral organ initiation, a process called floral meristem termination (FMT). FMT results in the loss of pluripotency of all the cells that remain in the FM, which will then be incorporated into production of the innermost organs of the flower.

Over the past few decades, studies using mitotic index and clonal sectors have revealed that most meristems are highly organized structures, composed of a central zone (CZ) and a peripheral zone (PZ), which harbor the stem cells and organogenic cells, respectively (Stewart & Dermen, 1970; Marc & Palmer, 1982; Steeves & Sussex, 1989). Thus, the maintenance of stem cell identity in the CZ is key to meristem indeterminacy, and many key genes controlling meristem homeostasis have been identified. Perhaps the most critical is the stem cell identity gene *WUSCHEL* (*WUS*), which is exclusively expressed at the base of the central zone (also called the organizing center (Laux *et al*., 1996; Mayer *et al*., 1998). The WUS protein functions non-cell autonomously to induce the expression of the gene *CLAVATA3* (*CLV3*), which codes for a small peptide that in turn diffuses down to repress *WUS* expression (Schoof *et al*., 2000; Yadav *et al*., 2011). This WUS-CLV3 feedback loop is critical for maintaining the homeostasis of stem cells in the meristems and was found to be widely conserved among land plants (Schoof *et al*., 2000; Lenhard, 2003; Yadav *et al*., 2011; Whitewoods *et al*., 2020). Therefore, when we consider the FMT process, we are explicitly asking when and how *WUS* expression is permanently down-regulated, a process that is best understood in the model systems *Arabidopsis thaliana* and tomato (Sun *et al*., 2009, 2014; Bollier *et al*., 2018). However, it remains to be seen whether these genetic mechanisms are more broadly conserved and, perhaps most importantly, both of these models have rather simple flowers with only four whorls of organs, representing only a small fraction of the diversity seen in angiosperms (Endress, 1990).

Another key component of investigating FMT from a developmental perspective is achieving a detailed understanding of cell division and expansion dynamics. Most previous work has relied on histology, scanning electron microscopy, *in situ* hybridization, or immunolocalization, all of which require fixation and, in most cases, sectioning of the tissues, rendering the dynamics static (Sappl & Heisler, 2013; Prunet & Duncan, 2020). Recent advancements in live imaging techniques have allowed researchers to analyze how gene activities regulate the spatial-temporal dynamic of cellular behaviors quantitatively, at unprecedented resolutions and in real time (Sappl & Heisler, 2013; Prunet & Duncan, 2020). Implementation of quantitative live imaging has provided answers to many long-standing questions in plant development, such as the timing of the establishment of adaxial/abaxial polarities during primordium initiation (Zhao & Traas, 2021), functions of the gene *SUPERMAN* in regulating organ boundaries and FMT (Prunet *et al*., 2017), and mechanisms generating the giant cells in the *A. thaliana* sepals (Roeder *et al*., 2010). Therefore, to gain an in-depth understanding of the dynamics of meristem function, we need both knowledge of different molecular regulatory networks and an understanding of how such networks directly control the precise actions of cell division and expansion over time, ideally in as many plant taxa as possible. So far, quantitative live imaging of meristems has only been developed for and applied to a small number of model species: the vegetative meristems of tomato, moss, and *A. thaliana* (Reddy & Roy-Chowdhury, 2009; Harrison *et al*., 2009; Hamant *et al*., 2019); the inflorescence meristems of *Gerbera hybrida, Brachypodium distachyon*, and *A. thaliana* (Heisler & Ohno, 2014; O’Connor, 2018; Zhang *et al*., 2021); the root meristems of *A. thaliana* (Rahni & Birnbaum, 2019); and the early FMs of *Cardamine hirsuta* and *A. thaliana* (Prunet *et al*., 2016; Monniaux *et al*., 2018).

In this study, we have applied a recently developed live confocal microscope imaging technique to produce the first quantitative characterization of the cellular dynamics in the FMs of *Aquilegia coerulea*, with a particular focus on FMT. *Aquilegia coerulea* is a member of the buttercup family (Ranunculaceae) and is a model system for evolutionary developmental studies with a well-annotated genome and a number of functional tools (Kramer, 2009; Filiault *et al*., 2018). We chose to focus on the developmental window of FMT because of its crucial role in flower development, but also because the cell dynamic changes during FMT have not yet been described quantitatively in any model systems. In addition, the FMs of *A. coerulea* have several significant differences compared to the FMs of *A. thaliana*, one being that they are maintained for a longer period before FMT to allow for the production of 15 to 17 whorls of floral organs compared to the four whorls of organs in *A. thaliana*. Moreover, FMT is very different between these two systems from a morphological perspective. After FMT, the carpel primordia of *A. thaliana* arise as a single syncarpous gynoecium (Hill & Lord, 1989), incorporating all of the cells that remained in the apex of the FM. By contrast, carpels of *A. coerulea* are formed from five distinct primordia (i.e., an apocarpous gynoecium) such that the apex of the FM is not spontaneously consumed by their emergence (Tucker & Hodges, 2005). Currently, we have no information regarding how different patterns of carpel primordia initiation influence FMT in flowering plants.

By imaging the same *A. coerulea* FMs in tissue culture at repeated time intervals and using the software MorphoGraphX (Barbier de Reuille *et al*., 2015), we conducted lineage tracing of cells and quantified the rate and distribution of cell division, as well as the degree and direction of cell expansion, over multiple developmental stages. Our results have revealed how the dynamics between cell proliferation and expansion change as the FM transitions from production of stamens to staminodes to carpels, allowing a detailed description of the initial developmental stages of floral organ primordia. To our knowledge, this study is the first quantitative live imaging of meristem development in a system with numerous whorls of floral organs and an apocarpous gynoecium and is the first cellular characterization of FMT. Our results will lay the foundation for investigating gene expressions in the *A. coerulea* FMs in real time, providing crucial information for our understanding of how the spatial-temporal regulation of meristem behavior is achieved in both an evolutionary and developmental context.

## RESULTS

### The developmental window covered in the study

The *A. coerulea* flower is composed of five organ types, which, arranged from outermost to innermost, are the sepals, petals, stamens, staminodes, and carpels (Fig. 1). The five sepals are initiated in spiral phyllotaxy while all other organs are produced synchronously in whorls of five (Fig. 1D). Each adjacent whorl initiates in alternating positions, either directly above the sepals or the petals, resulting in 10 orthostichies (i.e., vertical rows of organs) (Fig. 1C-E). Each flower consists of one whorl of sepals, one whorl of petals, an average of 10 to 12 whorls of stamens, two whorls of staminodes, and one whorl of carpels. Since we have focused on the later stages of FM development in the current study, the FMs we dissected for live imaging had typically initiated 8 to 10 whorls of stamens. For each stage, we designate the youngest primordial whorl as 1 and the previous whorl as 1+n; therefore, sd1 represents the inner whorl of staminodes that are produced after sd2, and st1 represents the last whorl of stamens initiated in the flower (Fig. 1E). In addition, we define “newly emerging primordia (NEP)” as the youngest organ primordia of the floral bud, which just bulges out at the periphery of the meristem and can be distinguished morphologically; while the “incipient primordia (ICP)” are the primordia that are initiating after the NEP but cannot yet be distinguished morphologically. Both NEP and ICP can be visualized by examining expression of the previously studied abaxial polarity gene *FILAMENTOUS FLOWER* (*AqFIL*) in *Aquilegia* (Fig. 1F; Meaders *et al*., 2020).

**Figure 1.**
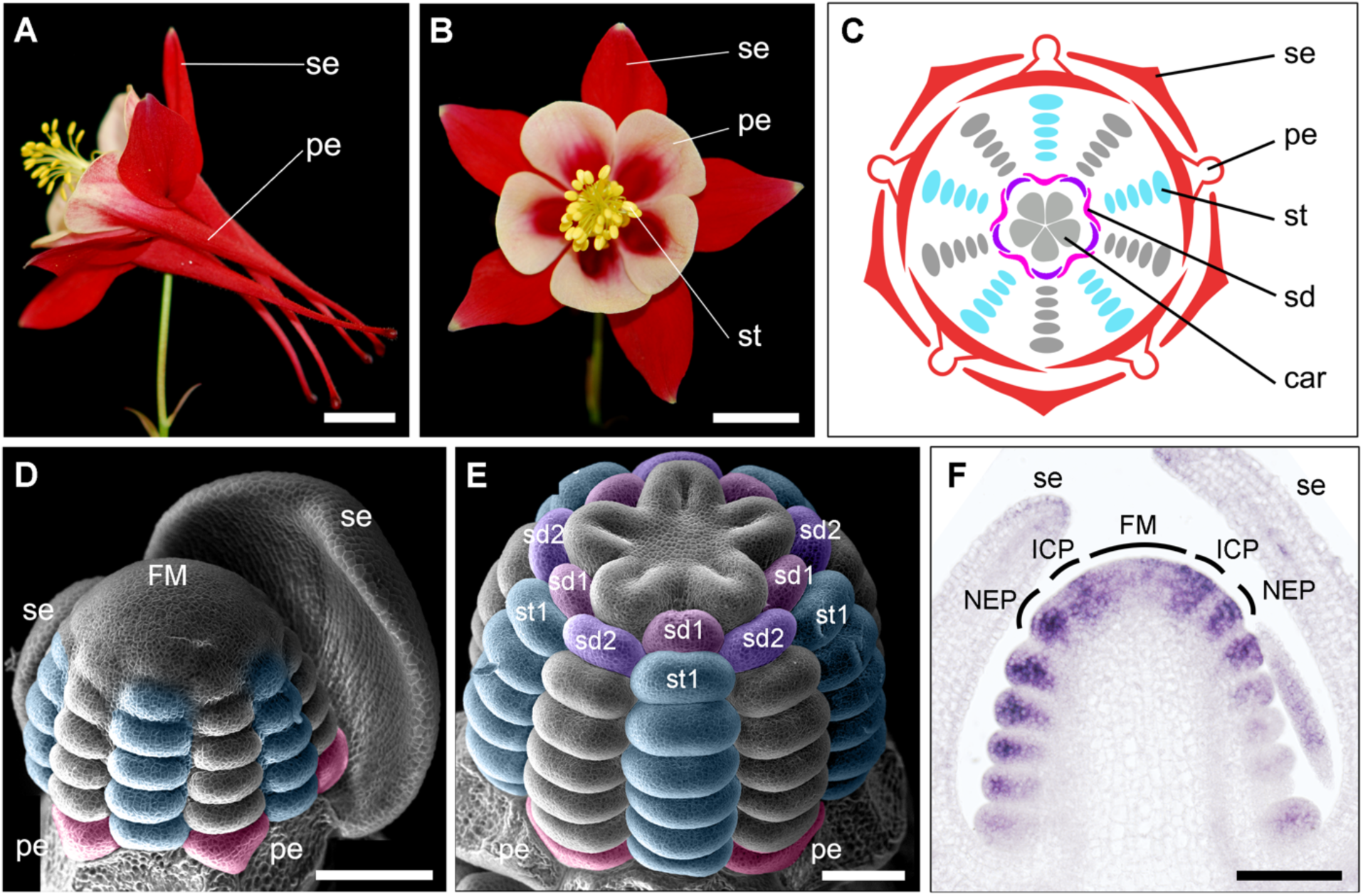
Floral morphology of *A. coerulea* flowers. (A) Side view of a mature flower. (B) Front view of a mature flower. (C) Floral diagram of a typical *A. coerulea* flower. (D) A young FM in the process of producing stamens. (E) A young floral bud in the initial stage of carpel development. (F) *In situ* hybridization of abaxial identity gene *AqFIL* in a young *A. coerulea* FM. Expression pattern was obtained by C. Meaders. se: sepals, pe: petals, st: stamens, sd: staminodes, car: carpels; FM: floral meristem; ICP: incipient primordia; NEP: newly initiated primordia. Scale bars: A, B = 1 cm; D-F = 100 μm.

We used a 48-hour imaging interval because we have observed that a new whorl of stamen or staminode primordia became visible roughly every 48 hours in our experimental conditions, and almost all cells that underwent a cell division during this imaging interval only divided once. Based on this imaging interval, we defined six time points (TP) in our study, resulting in five comparative developmental windows (Fig. 2). The first four TPs are distinguished by the successive initiation of floral organ whorls: st1 during TP1; sd2 during TP2; sd1 during TP3; and carpels during TP4. At TP5, the five carpel primordia formed a completely flat, star-shaped apex. At TP6, the carpel primordia each form an elevated ridge at their distal periphery. The meristematic dome was maintained until TP4, at which point the initiation of the carpel primordia resulted in flattening of the apex (Fig. 2). The overall distribution of observed cell sizes was consistent with previous observations (Steeves & Sussex, 1989) that cells in the center of the apex are generally larger than the cells in the periphery regions (Fig. 2).

**Figure 2.**
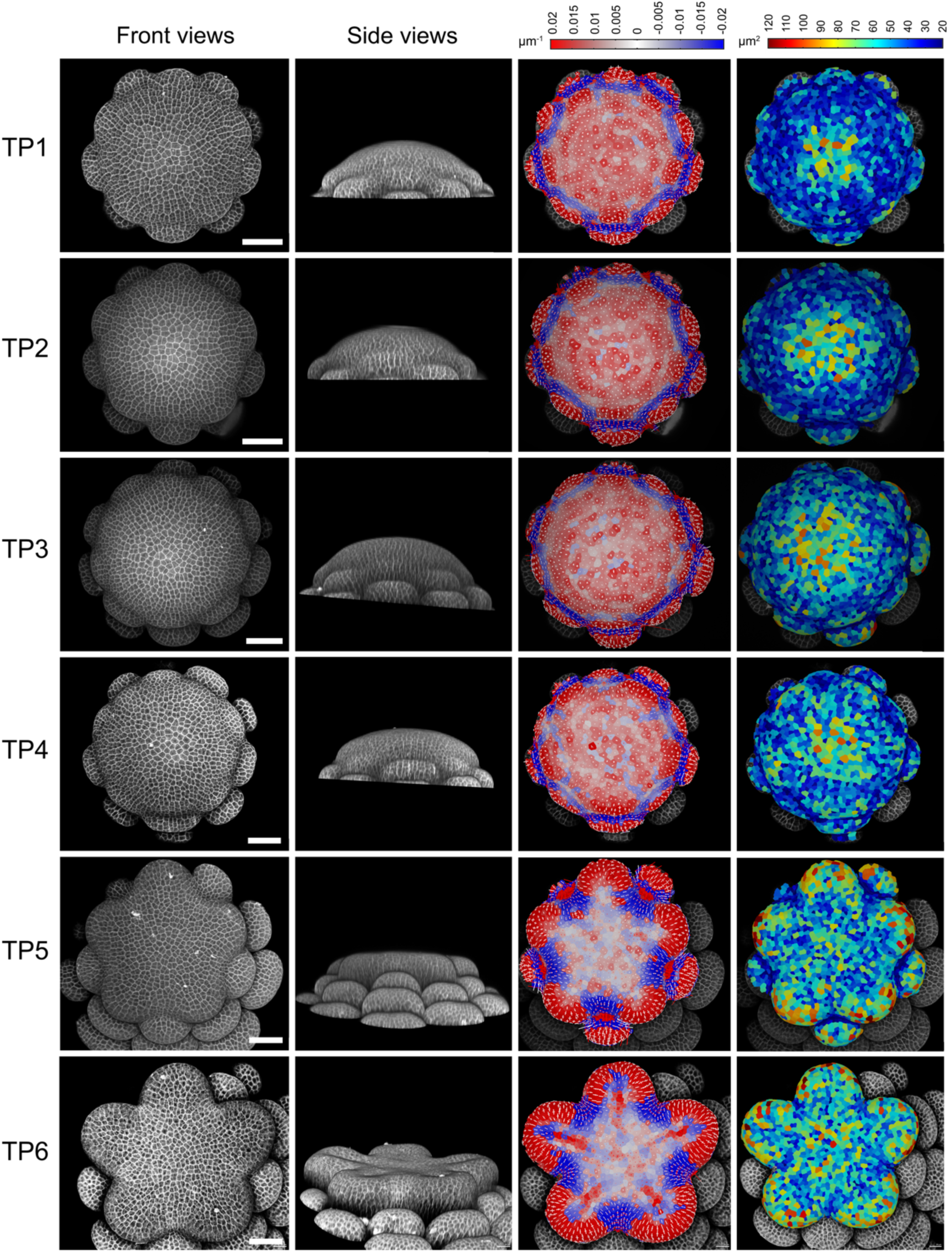
Developmental windows covered in the current study. Columns from the left to the right are the front view, side view, front view with curvature heatmap, and cell area heatmap of each time point 1-6 (TP1-6), respectively. Scale bars = 50 μm.

### Overall growth dynamics over the developmental windows

We used the imaging data to conduct lineage tracing of the epidermal cells of the floral apices to quantify their growth dynamics (Fig. 3). To determine whether there were any quantifiable differences in the patterns of growth between the TPs, we constructed growth alignment graphs by plotting the cell area expansion rates and the average number of cell divisions based on the locations of cells along the radial axis of each apex (Fig. 3, S1). For each apex, these radial transects were calculated from the center of the floral apex to the inner edges of the NEP whorl based on the curvature heatmap, dividing the transects into six equal bins (Fig. S1).

**Figure 3.**
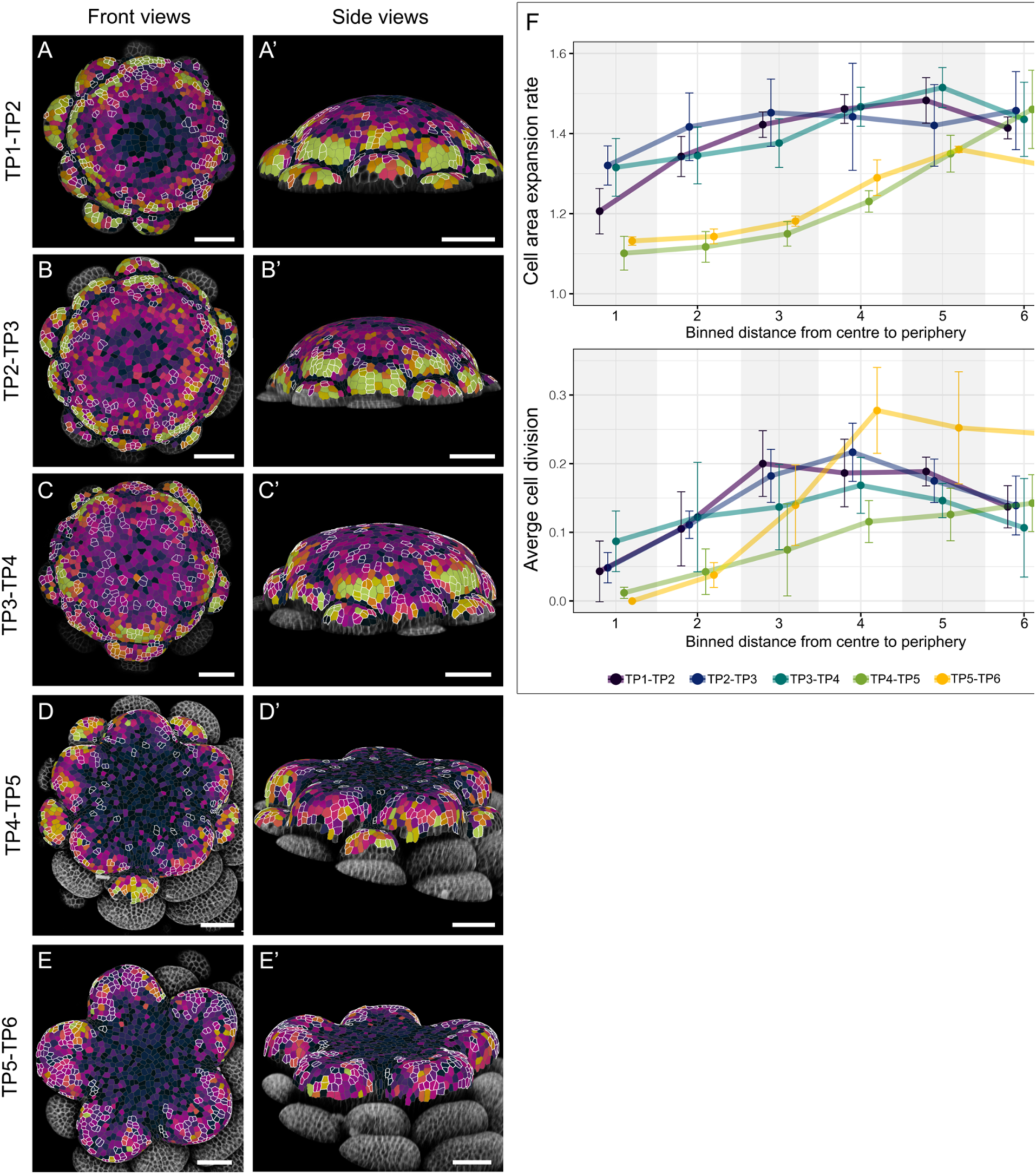
Overview of growth dynamics in each developmental window. (A-E’) Heatmaps showing the area expansion rates of all cells between two successive time points (TPs). If a cell has divided during the imaging interval, the sum of the cell areas of the daughter cells is compared to the area of the mother cell. All cells that experienced cell division during the interval are outlined in white. (F) Transects of cell growth from the center of the meristem to the most recently initiated primordia. These growth alignment graphs compare the cell area expansion rates (upper panel) and the average number of cell divisions (lower panel) between different developmental intervals. Error bars represented the standard error of biological replicates for each developmental interval. All developmental intervals had four biological replicates except for TP5-TP6, which had three biological replicates. Scale bars = 50 μm.

All developmental windows showed a general pattern that cell area expansion rates were lower at the center and higher at the periphery close to the NEP, but the overall expansion rates after TP4 (i.e., carpel primordia initiation; Fig. 3D-E’, F) were significantly lower than the earlier developmental windows (Table S1). The area expansion rates were relatively consistent across the entire meristem between TP2 to TP3 (Fig. 3B, B’, F), during which the sd1 primordia initiate. The cell area expansion rate displayed the largest difference between cells at the center and the periphery during the transition from TP4 to TP5 (Fig. 3D, D’, F), when the newly initiated carpel primordia were forming the flat star apex. Interestingly, the TP5 to TP6 (Fig. 3E, E’) window showed very similar cell area expansion patterns compared to TP4 to TP5 (Fig. 3D, D’), only differing in cells close to the periphery, in which the cell area expansion rates are significantly higher in TP4 to TP5 (Fig, 3F; Table S1).

Overall, rates of cell division were uniformly low in the center of the meristems across all time points (Fig. 3D-E’, F), although there was a significant uptick between TP3-TP4 preceding initiation of the carpel primordia (Fig. 3C, C’; Table S1). Moving outward in the meristems, rates of cell division generally increased progressively, regardless of the stage (Fig. 3). The one distinct exception was in TP5-TP6, during which rates were dramatically low in the center but sharply increased in the third bin domain, a pattern that was statistically different from the other stages (Fig. 3F; Table S1).

### Floral organ primordia initiation

Subsequently, we analyzed how the initial outgrowth of organ primordia was achieved. Using st1 primordium initiation as an example, at TP1 the physical bulging of the primordia became visible (Fig. 4A, D, G). To precisely describe the growth each primordium, we defined the lateral axis as the axis that is parallel to the circumference of the dome of the FM, and the radial axis as the axis that runs from the center of the meristem to the center of the primordium (Fig. 4A). At TP1, Fig. 4A shows 10 cells of this NEP that are outlined in blue, representing the majority of the cells that exhibited positive surface curvature (i.e., are physically bulging out) compared to their surrounding cells, while the 10 cells outlined in green mainly had negative surface curvature (Fig. 4D). This includes part of the boundary region that separates this NEP from the primordium in the whorl positioned below (Fig. 4D). At TP2, the cells of the green domain, which have become the abaxial surface of the primordium (Fig. 4E), were the only cells that exhibited high growth rates during this developmental window (Fig. 4H). Cell division occurred on both the abaxial and adaxial sides (Fig. 4H), suggesting that this higher growth rate was mainly driven by cell expansion, which was supported by the cell anisotropy map (Fig. 4J). Cells on the abaxial side exhibited strong anisotropy along the radial axis, while cells on the adaxial side almost exclusively showed cell expansion along the lateral axis (Fig. 4J). At TP3, cell division occurred on both adaxial and abaxial sides of the primordium, and almost all cells in the primordium were exhibiting relatively high growth rates (Fig. 4I). The direction of cell expansion for the abaxial cells were more isotropic at this point (Fig. 4K). Interestingly, the adaxial-most layer of the cells at TP3 experienced significant compression along the radial axis while continue to expand along the lateral axis (Fig. 4K), indicating these cells were incorporated into the formation of the organ boundary.

**Figure 4.**
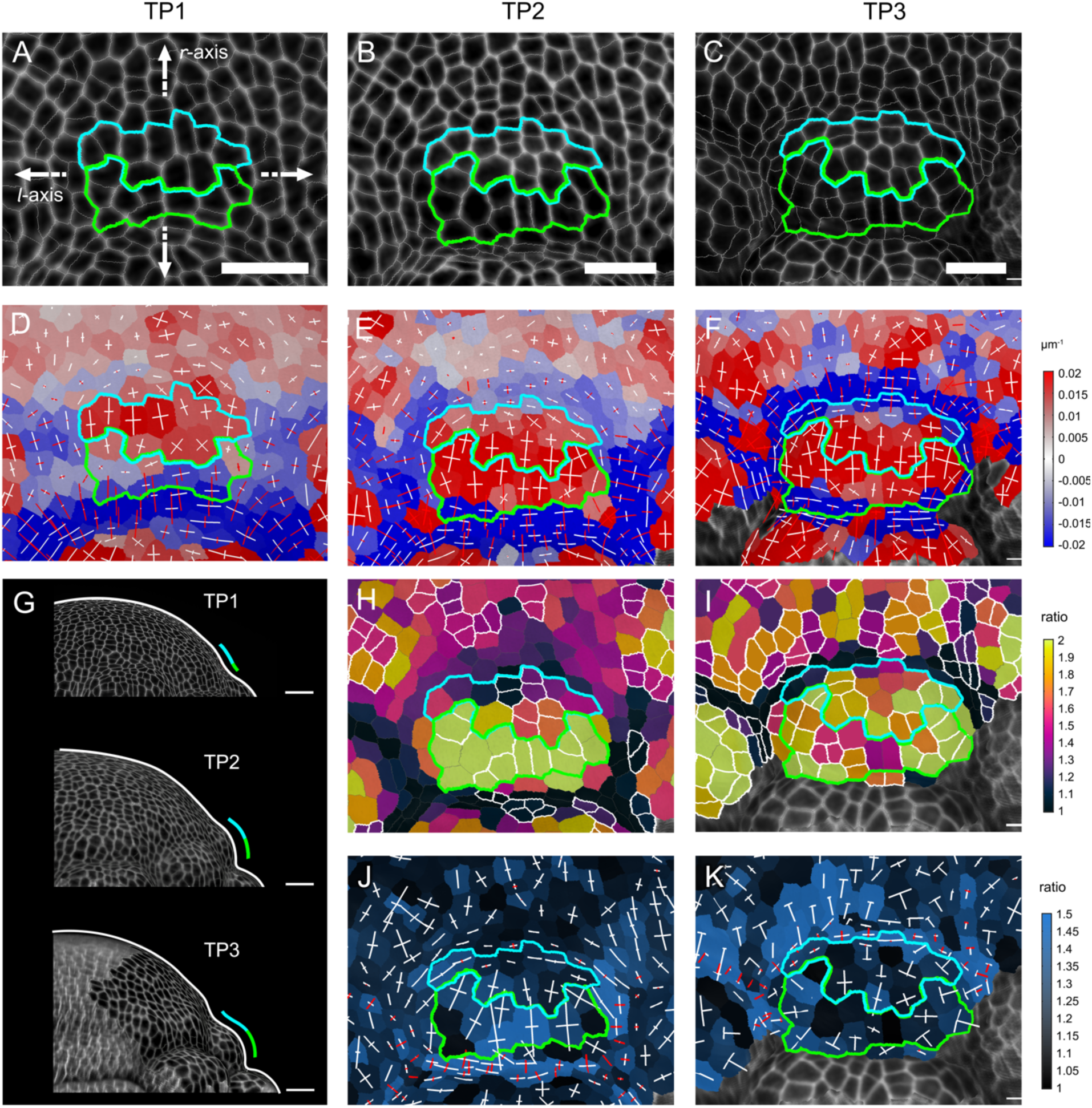
Initiation of st1 primordia. (A-C) Front view of a st1 primordium at TP1, TP2, and TP3, respectively. Cells outlined in green show the highest growth rates at TP1-TP2 (H) while cells outlined in blue show the highest curvature at TP1 (D). (D-E) Surface curvature heatmaps of A, B, and C, respectively. (G) Side views of the FM at the equivalent TPs. The white line outlines the overall shape of the FMs while the blue/green lines indicate the side view of the st1 primordium. (H, I) The cell area expansion heatmaps between TP1 to TP2, and TP2 to TP3, overlaid on B and C, respectively. Cells outlined in white are cells that experienced cell division during the interval. (J, K) Cell expansion anisotropy heatmaps between TP1 to TP2, and TP2 to TP3, overlaid on B and C, respectively. The color of a cell represents the value of the ratio between the changes in the longest axis and the shortest axis during the interval, and lines inside of the cell represent the degree and direction of cell expansion. White and red lines show the expansion, or compression, of the cell shape along the indicated axis, respectively. *r-*axis: radial axis; *l-*axis: lateral axis. Scale bars: A, B, C = 20 μm; G = 50 μm.

This phenomenon, that the initial outgrowth of the primordium was mainly achieved by high growth rates of the abaxial cells, was also observed for the initiation of staminode and carpel primordia (Fig. 5). Before entering the rapid growth phase, the abaxial cells were initially comprised of only one or two layers and were less than 10 cells wide (e.g., sd1 in Fig. 5A and car in Fig. 5B). These cells were located at the abaxial-most position of the ICP, and many (if not most) of them had negative surface curvatures (e.g., sd1 in Fig. 5D and car in Fig. 5E). They then exhibited substantial, directional cell expansion along the radial axis, coupled with cell divisions so that they quickly made up half of the primordium (e.g., sd1 in Fig. 5H, J and car in Fig. 5I, K). Subsequently, the cells on the adaxial side started to exhibit relatively strong growth, and the direction of expansion of most of the cells in the primordium became more isotropic (e.g., sd1 in Fig. 5K).

**Figure 5.**
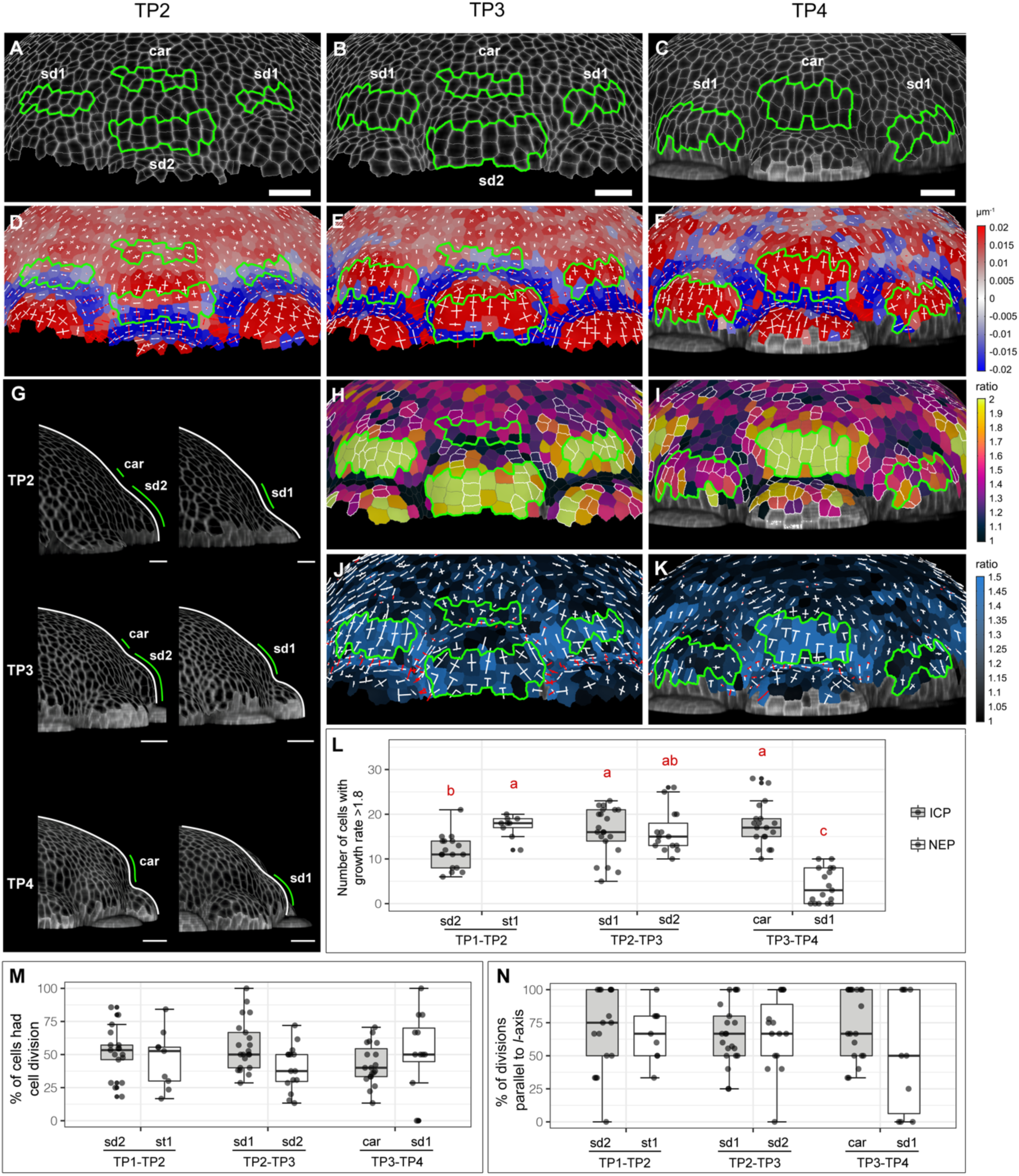
Initiation of staminode and carpel primordia. (A-C) Front view of the sd1, sd2, and car primordia at TP2, TP3, and TP4, respectively. The cells outlined in green are the abaxial cells of interest of the initiating primordia, as defined by high growth rates during TP2-TP3 (H) or TP3-TP4 (I). (D-E) Surface curvature heatmaps of A, B, and C, respectively. (G) Side views of the FM at different TPs. The white line outlines the overall morphology of the FMs and the green line indicates the position of the primordia of interest. (H, I) Cell area expansion heatmaps between TP2 to TP3, and TP3 to TP4, overlaid on B and C, respectively. Cells outlined in white resulted from cell division during the interval. (J, K) Cell expansion anisotropy heatmap between TP2 to TP3, and TP3 to TP4, overlaid on B and C, respectively. The color of a cell represents the value of the ratio between the changes in the (Figure 5 continued) longest axis and the shortest axis during the interval, and lines inside of the cell represent the degree and direction of cell expansion. White and red lines show the expansion, or compression, of the cell shape along the indicated axis, respectively. (L) Comparisons of the number of cells with growth rates > 1.8 in primordia of different stages and developmental windows. Different letters above the boxplots represent significant differences between different primordia (p < 0.05, using ANOVA followed by Tukey’s pairwise multiple comparisons). (M) Comparisons of percentages of cells that had growth rates > 1.8 and also experienced cell division during different time points. No significant difference was detected between different primordia using ANOVA. (N) Comparisons of the orientation of cell division among the cells that both had growth rates >1.8 and experienced cell divisions. No significant difference was detected between different primordia using ANOVA. For (L-M), each data point is a primordium, and for each developmental window, at least three FMs were quantified. Note that for each time period, the identity of the ICP and NEP shift. For instance, sd2 are the ICP at TP1-TP2 but become the NEP at TP2-TP3. Scale bars: A, B, C = 20 μm; G = 50 μm.

In addition, we observed that after the initial strong growth phase in the abaxial cells, the primordia of sd1 displayed a lower overall growth rate than other NEP; for instance, sd1 in Fig. 5H vs. sd2 in Fig. 5I. This observation was confirmed by quantifying the number of cells that exhibited growth rates above 1.8 in ICP and NEP of different developmental windows (Fig. 5L). During the developmental window TP2 to TP3, in which the sd1 was the ICP, these primordia had similar numbers of cells exhibiting high growth compared to ICP and NEP of other stages. However, once the floral buds enter the TP3 to TP4 window, in which the carpels were the ICP and sd1 was the NEP, the number of sd1 cells exhibiting high growth decreased significantly (Fig. 5L).

Among the cells that exhibited high growth rates in the ICP and NEP of different developmental windows, we asked what percentage of these cells experienced cell division during the interval. We did not observe any significant difference between primordia of different windows (Fig. 5M) and, on average, about 50% of the cells exhibited both high growth and experienced cell division. We also examined the orientations of the division planes among these cells, and found that while there was no significant difference between primordia of different stages and developmental windows, most of the cell divisions occurred parallel to the lateral axis (Fig. 5N).

### Early carpel development

We then analyzed the initial development of the carpel primordia, with a particular focus on three diagnostic regions: the distal edge, the primordial boundary, and the adaxial fold (Fig. 6A). From TP4 to TP6, the width of the distal edge increased both along the lateral axis and the radial axis, and the curvature of most cells increased (Fig. 6B-D). Over the same period, the primordial boundary became more folded towards the center of the apex (Fig. 6E-G). At TP4 and TP5, the regions that would become the adaxial folds transition from having positive surface curvature to having close to zero (i.e. flat) surface curvature (Fig. 6H, I). At TP6, invagination of the adaxial fold appeared to initiate at the base of the primordium and propagate outward such that the proximal region was more deeply folded than the distal at this stage (Fig. 6J).

**Figure 6.**
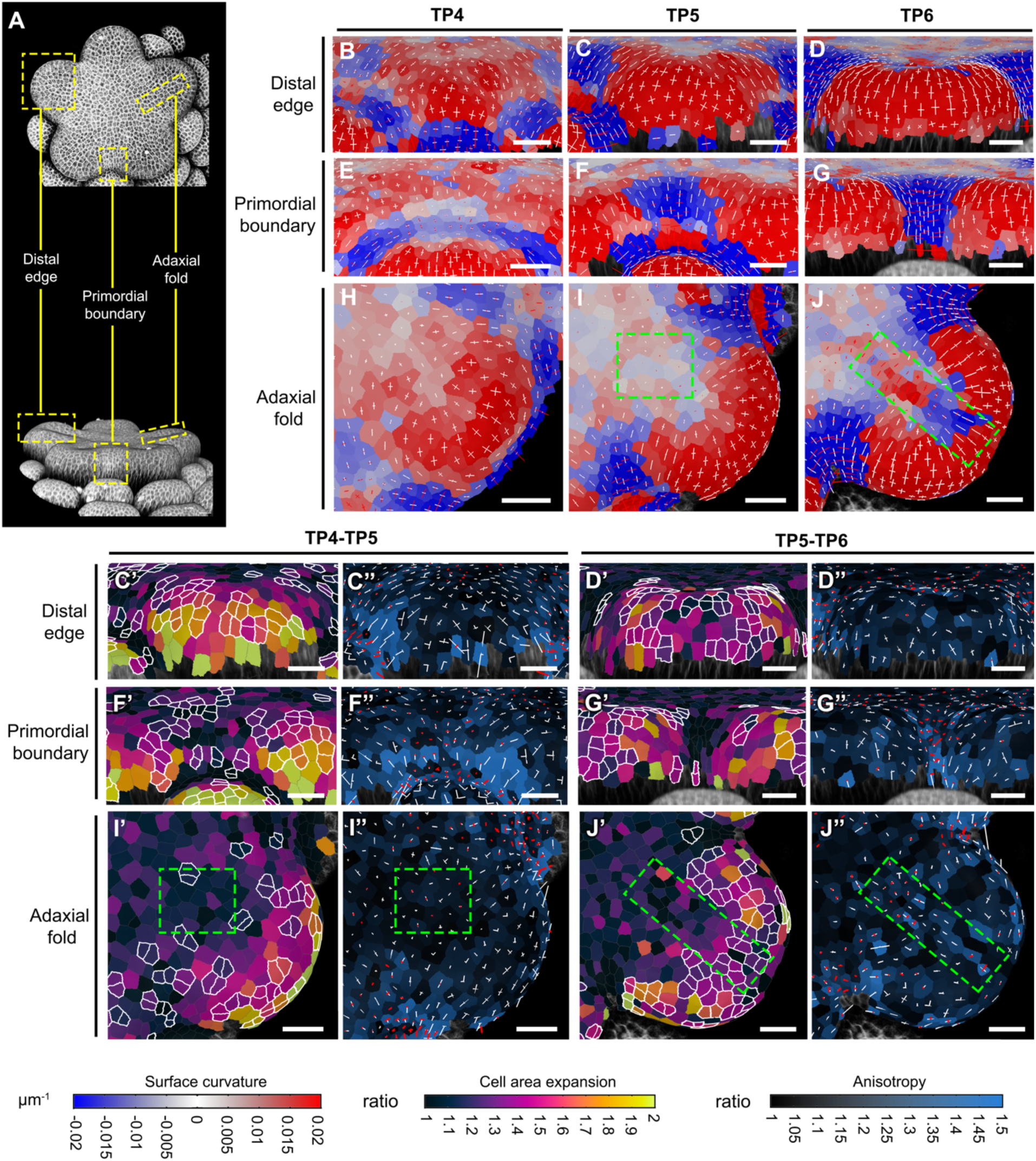
Early carpel primordia development. (A) Front view and side view of an *A. coerulea* FM at TP6 to indicate different regions of interest. (B-D) Side views of the surface curvature heatmaps of the distal edges of carpel primordia at TP4, TP5, TP6, respectively. (E-G) Side views of the surface curvature heatmaps of the primordial boundaries between carpels at TP4, TP5, TP6, respectively. (H-J) Front view of the surface curvature heatmaps of the adaxial folds of carpel primordia at TP4, TP5, TP6, respectively. (C’, C’’) Heatmaps of cell area expansion and (Figure 6 continued) anisotropy of distal edges between TP4 to TP5, respectively, overlaid onto TP5. (D’, D’’) Heatmaps of cell area expansion and anisotropy of distal edges between TP5 to TP6, respectively, overlaid onto TP6. (F’, F’’) Heatmaps of cell area expansion and anisotropy of primordial boundaries between TP4 to TP5, respectively, overlaid onto TP5. (G’, G’’) Heatmaps of cell area expansion and anisotropy of primordial boundaries between TP5 to TP6, respectively, overlaid onto TP6. (I’, I’’) Heatmaps of cell area expansion and anisotropy of adaxial folds between TP4 to TP5, respectively, overlaid onto TP5. (J’, J’’) Heatmaps of cell area expansion and anisotropy of adaxial folds between TP5 to TP6, respectively, overlaid onto TP6. Cells outlined in white in all cell area expansion heatmaps were cells that experienced cell divisions during the intervals. Green dashed boxes in I, I’, I’’, J, J’, J’’ indicate the adaxial folds. All distal edge panels are the side view of the distal edge, all primordia boundary panels are the side views of the primordial boundary, and all adaxial fold panels re the front view of the adaxial fold. Scale bars = 20 μm.

Once the five carpel primordia were initiated, two processes happen simultaneously to achieve the flat star shape (Fig. 6). First, cells at the distal edges of the carpel primordia continued to display relatively high growth rates (Fig. 6C’), while cells at the center of the floral apex exhibited much lower overall growth rates (Fig. 3). This had the effect of elevating the carpel primordia to the same level as the center of the floral apex. However, unlike the early initiation of the carpel primordia, during which growth was mainly driven by the strong anisotropic expansion from the abaxial cells (Fig. 5I, K), the overall anisotropy of the cells on the carpel primordia distal edges was lower and less directional (Fig. 6C”). Meanwhile, numerous cells underwent cell division during this developmental window, suggesting that growth was mainly driven by cell division (Fig. 6C’). Second, cells at the primordial boundaries experienced strong compression along the lateral axis (Fig. 6F”), which helped to define the primordial boundary regions of the flat star shape. Interestingly, at this stage, although the floral apex was flat and the adaxial folds were not yet physically visible (Fig. 6I), minor compression of cells in the future location of the adaxial fold can already be observed (Fig. 6I”).

After formation of the flat star apex, the carpel primordia began to elevate along their distal ridges, which was mainly achieved by concentrated cell divisions on the ridges (Fig. D’, J’) while the anisotropic expansion rates of cells on the ridges remained low (Fig. 6D”). The growth alignment graphs in Fig. 3 also support this observation of concentrated cell divisions in the peripheral region relative to the earlier developmental windows (Fig. 3). Meanwhile, the cells at the primordial boundaries continued to experience strong compression along their lateral axis. We did not observe any further cell division in the primordial boundaries (Fig. 6G’, G”). In addition, the adaxial folds of the carpels have become morphologically visible (Fig. 6J), exclusively achieved by modifications in cell shape since no cell division occurred in the region (Fig. 6J’). Specifically, cells of the adaxial fold located closer to the center of the floral apex experienced strong compression, while those in the distal region of the adaxial fold exhibited strong anisotropic expansion along the radial axis (Fig. 6J”).

## DISCUSSION

### Cellular dynamic change during FMT in *A. coerulea*

Despite its essential role in floral development, little is known about meristem dynamics during FMT at a cellular level, and the molecular basis of FMT has only been investigated in *A. thaliana* and tomato (Sun *et al*., 2009, 2014; Bollier *et al*., 2018). In *A. thaliana*, FMT is marked by down-regulation of the gene *WUS*, which determines stem cell identity; this shift also coincides with the initiation of carpel primordia (Sun *et al*., 2009, 2014). Therefore, FMT defines the transition from being pluripotent to organogenic for the cells in the central zone, and the precise timing of *WUS* repression is considered a key factor that determines the number of cells produced for carpel development (Sun *et al*., 2014; Sun & Ito, 2015).

Previous transcriptomic sequencing in young *A. coerulea* FMs indicated that the expression of *AqWUS* is maintained during the developmental stages equivalent to TP1 through TP3 in the current study, with expression dropping rapidly to undetectable levels during the developmental stages equivalent to TP4 to TP6 (Min & Kramer, 2020). If we examine the central region of the FM (i.e., bin1 and bin2 in the growth alignment graphs, Fig. 3F), which roughly corresponds to the central zone, we did observe similarly low numbers of cell divisions between TP1 to TP3 (Fig. 3F; Table S1), with bin2 being higher than bin1. During TP4 to TP6, which corresponds to the early stages of carpel development, the average number of cell divisions in the center of the floral apex was close to zero (Fig. 3F; Table S1). However, most strikingly, we observed a higher number of cell divisions in the center of the floral apex between TP3 to TP4, during which the carpel primordia are initiated (Fig. 3C, C’, F), with the bin1 values being statistically significantly higher than all other stages (Fig. 3F; Table S1).

This observation raises some intriguing questions: Why is there an increase in cell division in the central zone when the carpel primordia are initiating, and what is responsible for the pattern? This period also appears to correspond with the onset of FMT based on the decline in *AqWUS* expression, so is this the last flush of CZ cell divisions or are the divisions related more directly to initiation of the carpel primordia? Answering these questions will require a better understanding of how *AqWUS* down-regulation is controlled in *Aquilegia*, particularly the nature of any “timer mechanism” regulating the number of cell divisions before FMT. In the FMs of *A. thaliana* and tomato, the gene *KNUCKLES* (*KNU*) is activated in a cell division-dependent manner, which determines the temporal control of FMT, since once *KNU* is activated, it terminates the expression of *WUS*. Manipulation of cell cycles in the FM can accelerate or delay the activation of *KNU*, which results in the premature or delayed termination of the FM, respectively (Sun *et al*., 2014). We currently do not know whether the *KNU*-*WUS* pathway is conserved in other angiosperm systems, especially taxa with more than one whorl of stamens and unfused carpels but, even if it is not conserved, it will be interesting to examine whether a division-dependent timing mechanism is used for FMT in general. Overall, the combined patterns of cell behavior we observe at the center of the meristem during TP3-TP4 then transitioning into TP4-TP5, which include a sudden uptick in divisions during TP3-TP4 and a sharp decline in cell expansion following this phase, appear to be markers for a loss of stem cell identity in this zone in conjunction with the FMT.

### Floral organ primordia initiation in *A. coerulea*

In following the initiation of stamen, staminode, and carpel primordia, we observed interesting dynamics in the relative growth behavior of the adaxial and abaxial cells (Fig. 4; Fig. 5). First, only some of the cells that initially exhibited positive curvature in the NEP end up as adaxial cells in the primordium proper, with the adaxial-most cells being incorporated into the primordium boundary. Second, cells with negative surface curvature that were mainly located in the boundary region below the initial bulge ultimately become the cells exhibiting the highest growth rates. These high growth rates were primarily driven by highly anisotropic cell expansion, which coupled with later cell division quickly promotes outgrowth of the abaxial surface of the primordium. This pattern is consistent with a previous study of expression of the abaxial polarity gene *AqFIL*, which is detected broadly across the entire ICP and most of the NEP (Fig. 1F), similar to the expression patterns of *FIL* in *A. thaliana* FMs (Meaders *et al*., 2020; Zhao & Traas, 2021). It is curious that cells exhibiting the highest growth rates are along the extreme abaxial margin of the primordium, representing only a portion of the typical abaxial domain, a pattern that has also been observed during leaf and sepal primordium initiation in *A. thaliana* (Zhao & Traas, 2021). A close examination of the spatial-temporal expression patterns of other polarity genes in *A. coerulea* will be a good starting point to decipher any potential difference in the abaxial cells, as well as the transition of cells that were originally in the adaxial surface but ended up in the organ boundary region.

The observed difference in the growth dynamics between sd1 and sd2 was more surprising. Once the primordia initiate as NEP, the growth rates of sd1 appeared to be lower than other NEP and ICP in other developmental intervals (Fig. 5L). Although the sd1 and sd2 whorls have the same staminode identity (Kramer *et al*., 2007; Sharma & Kramer, 2013), a previous study has observed subtle morphological differences that are apparent when the primordia are as small as 1-2 mm in length, most notably lateral marginal curling that facilitates late-stage adhesion between the organs (Meaders *et al*., 2020). In that study, the authors raised the question of whether these differences were developmentally determined from the earliest stages or were due to some kind inductive interaction of the tissues. Our observation of differences from inception may suggest that they do harbor distinct developmental trajectories from their earliest stages, possibly implicating identity-based differences.

In addition, we examined the early developmental processes of the carpel primordia in detail, capturing a transition from growth that is mainly driven by strong anisotropic cell expansion to one that is promoted by concentrated cell divisions (Fig. 6). The initial cell expansion phase helps to lift the carpel primordia up, while the cell division phase appears to sculpt their shape (Fig. 5I, K; Fig. 6). The first phase of cell expansion seems to be very similar to the initiation of the stamen and staminodes, which was likewise driven by strong anisotropic expansion by a small number of abaxial cells (Fig. 5). However, the second phase of carpel growth is quite different than the other organs in that there is a much stronger reliance on accumulation of cell divisions while the cell areas did not expand significantly (e.g., st1 in Fig. 4 compared to Fig. 6). This observation could suggest that we observed a transition between two sets of molecular programs during early carpel primordia growth: one for the earliest phase of primordia initiation that is common to all floral organs and the subsequent program that is specific to sculpting carpel primordia.

## MATERIALS AND METHODS

### Plant materials, growth conditions, and dissection

Seeds of *Aquilegia* x *coerulea* ‘Kiragami’ were purchased from Swallowtail Garden Seeds (Santa Rosa, CA, USA) and germinated in damp soil. The regular growth condition for seedlings and young plants is 16 h daylight at 18 °C and 8 h dark at 13 °C, with humidity under 40%. Once the plants developed approx. six true leaves, they were transferred into vernalization conditions (16 h daylight at 6 °C and 8 h dark at 6 °C) for three to four weeks, and then moved back to the regular growth conditions. When the primary inflorescences started to develop, young side branches with axillary meristems were cut off from the plant, washed in 10% bleach for 20 min, and then thoroughly rinsed with double-distilled water three times. Axillary meristems on the branches were then dissected using a surgical needle under a dissecting microscope. After all the sepals were removed, the floral meristems placed on a petri dish with tissue culture medium composed of 0.5X Linsmaier & Skoog medium (Caisson Labs, Smithfield, UT), with 3% sucrose, 0.8% UltraPure Agarose (Invitrogen), 10^−6^ M kinetin (MilliporeSigma, Burlington, MA), and 10^−7^ M gibberellic acid (GA3, MilliporeSigma, Burlington, MA). The Petri dishes were placed in a tissue culture growth chamber with 16 h light at 20 °C and 8 h dark at 13 °C.

### Imaging

FMs were stained with 0.5mg/mL propidium iodide solution in double-distilled water (Sigma-Aldrich, Inc, St. Louis, MO) by immersing the tissue in stain solution for 2.5 minutes for the initial timepoint and then 2 minutes for subsequent timepoints. The stain was removed, and the tissue washed in double distilled water three times. Meristems were imaged immediately after staining using a LSM 980 NLO Multi-photon confocal laser scanning microscope (Ziess, Germany) equipped with a water immersion objective (W Plan-Apochromat 20x/1.0 DIC UV-IR M27 75mm, Ziess). A DPSS 514nm laser was used for excitation and emission was collected between 580-670nm. Scans were frame averaged 2x and z-sections taken at 2μm intervals. After imaging, the remaining water in the petri dishes was carefully removed using a pipette and the petri dishes were returned to the issue culture growth chamber. All samples were imaged every 48 hours, and most of the samples were imagined up to three to four times (i.e., 3 to 4 TPs). Since in the current study we have covered a developmental window covered 6 TPs in total, we have stacked multiple independent time-lapse imaging series to cover the 6 TPs. For instance, there are four independent time-lapse imaging from spanning stages that were equivalent from TP1 to TP4, and four independent time-lapse imaging spanning stages that were equivalent from TP3 to TP6. Every successive TP interval had at least three biological replicates (i.e., independent time-lapse imaging sessions), which were also used as biological replicates to construct the growth alignment graphs (Fig. 3F). Stacking several time-lapse experiments (with replicates) to achieve a larger developmental window is common in long-time live imaging studies (e.g. Kuchen *et al*., 2012; Kierzkowski *et al*., 2019).

### Imaging processing and data analysis

The confocal image files were transformed into .tif files in ImageJ (https://imagej.nih.gov/ij/) and processed in MorphoGraphX (https://morphographx.org/) following the steps listed in https://kramerlab.oeb.harvard.edu/protocols. Briefly, all image stacks were Gaussian Blurred twice with a radius of 1 on all X/Y/Z sigma, and all meshes were subdivided and smoothed (with 10 passes for each smooth) three times. Cells on the surfaces were segmented by using auto-segmentation, and all the segmentation errors were corrected manually. Heatmap graphs and values were generated by MorphoGraphX, and statistical analysis (ANOVA, Tukey’s HSD) was done in R (version 1.1.456). For the growth alignment graphs, we had four biological replicates for every developmental interval except TP5-TP6, which had three biological replicates. The distance between the center of each floral apex to the edge of the newly emerged primordia (NEP; defined in Results) was normalized and divided into six equal bins. For each bin, the biological replicates were pooled and means for cell area expansion and cell division were calculated.

## AUTHOR CONTRIBUTIONS

YM and EMK conceived of the study. YM and SJC imaging, data processing and analysis, and EMK provided oversight of the study. YM wrote the manuscript with input from SJC and EMK.

## ACKNOWLEDGEMENT

The authors would like to thank Mingyuan Zhu, Daniel Kierzkowski, Anne-Lise Routier-Kierzkowska, Richard Smith, and Soeren Strauss for patiently answering all the questions about MorphoGraphX, and Weilin Meng for generously lending his PC during the entire Covid19 pandemic, on which we conducted all the image processing and analysis. This study was funded by the Emerging Models Award to Ya Min from the Society of Developmental Biology.

## Supplemental materials

### List of files

**Figure S1.**
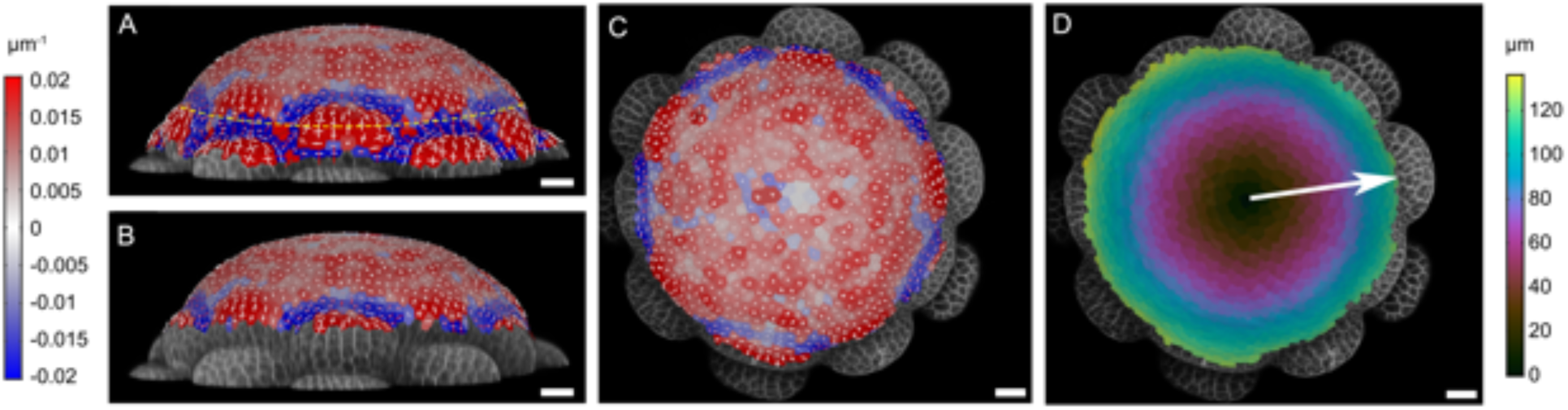
Regions used for constructing growth alignment graphs. (A) Side view of a meristem overlayed with surface curvature heatmap. Dashed yellow line circled the region defined by the boundary of the ITP. (B) Side view of a meristem overlayed with surface curvature heatmap only with the regions that will be used to construct the growth alignment graphs. (C) Front view of (B). (D) Cell distance heatmap that was used to determine the center most cell of the region of interest, and the white arrow represented the axis that would be used to construct the growth alignment graphs. Scale bars = 20 μm.

**Table S1.**
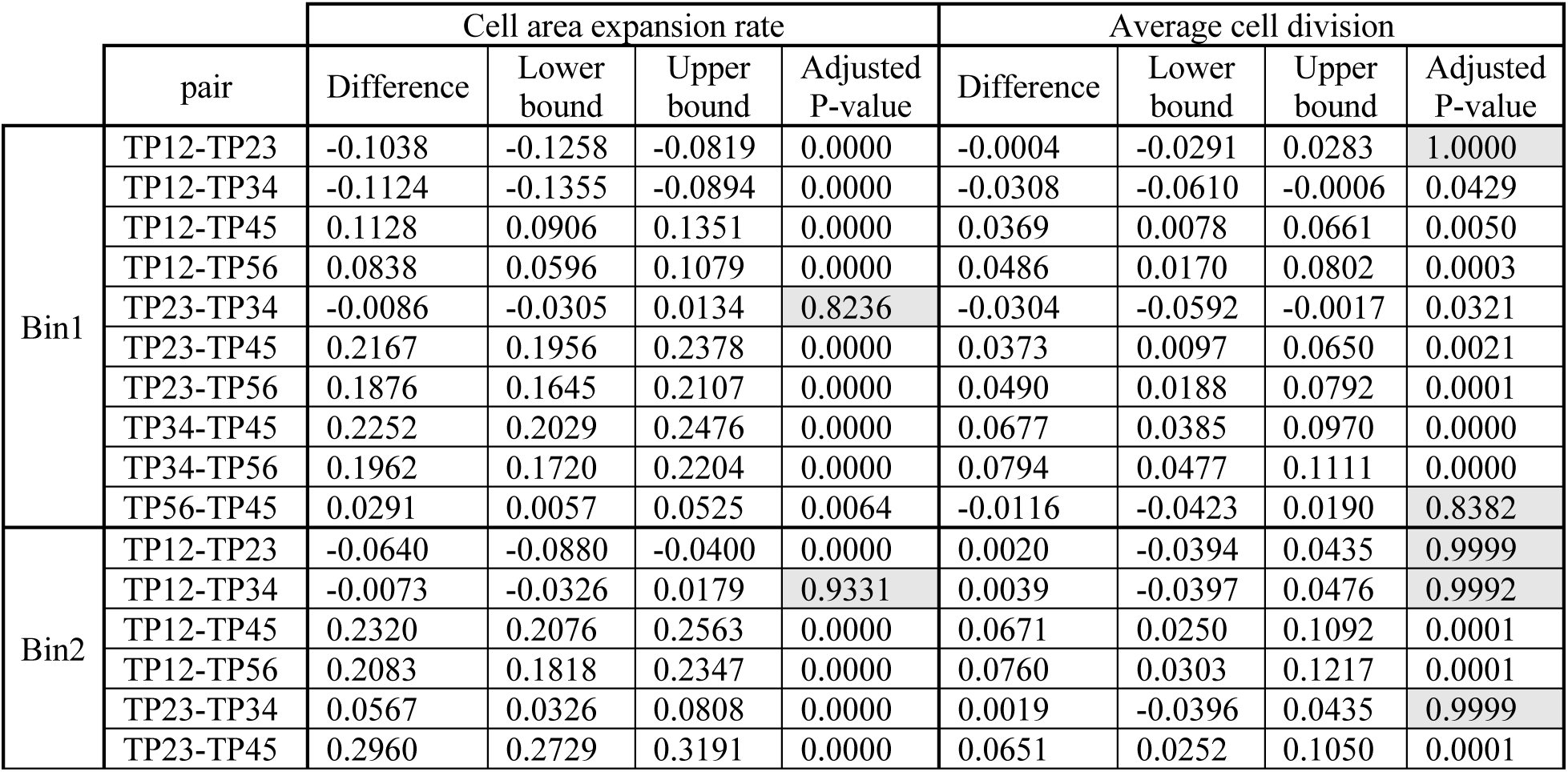
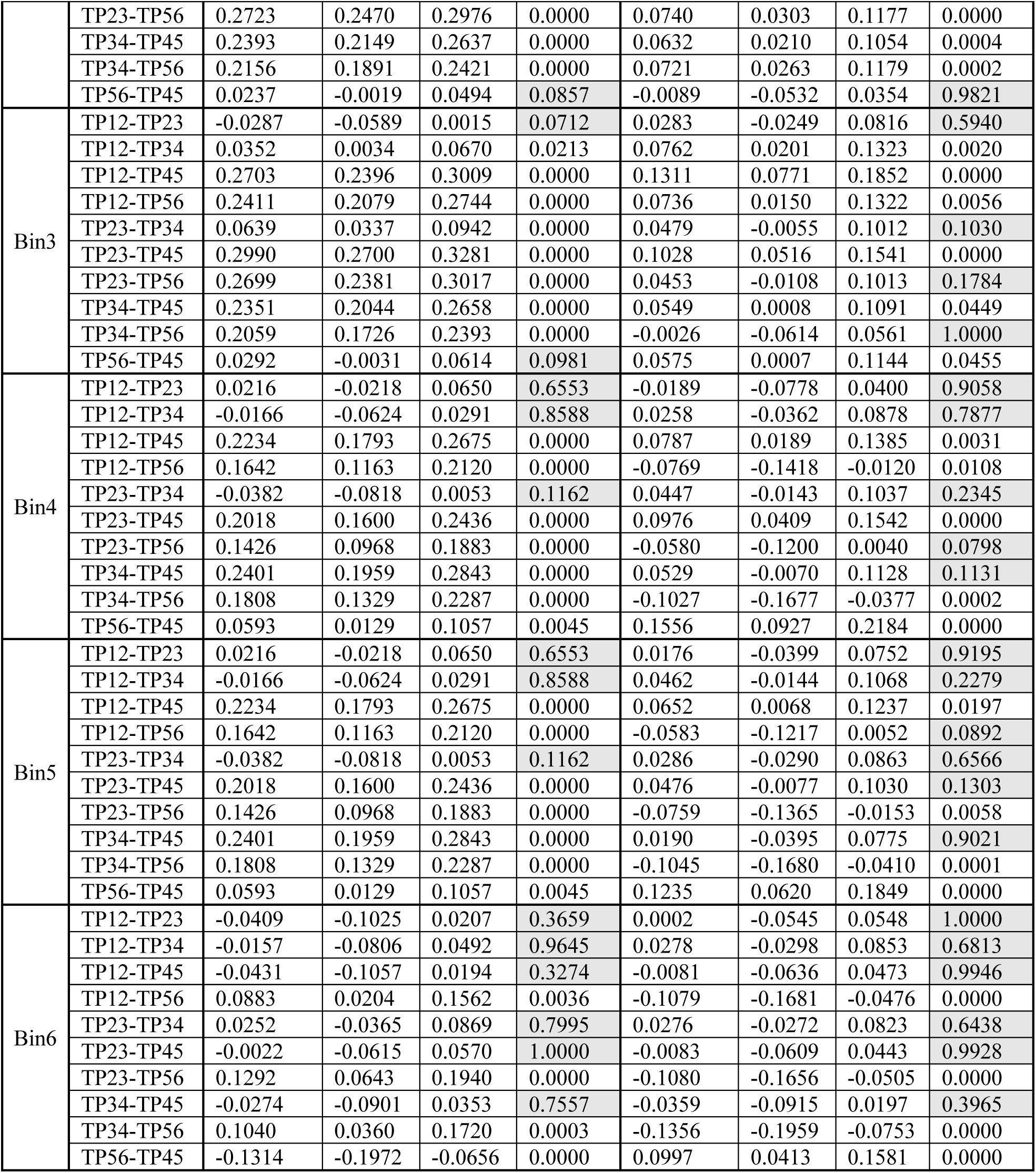
Pair-wise Tukey’s HSD test for cell area expansion rates and average cell division numbers between different TP intervals for the growth alignment graphs. Each interval is abbreviated as TP(first TP)(second TP): e.g. TP12 stands for the interval between TP1 to TP2. Cells in grey were pairs that have no significant difference.

